# Mutations upstream from *sdaC* and *malT* in *Escherichia coli* uncover a complex interplay between the cAMP receptor protein and different sigma factors

**DOI:** 10.1101/2023.10.26.564124

**Authors:** Pernille Ott Frendorf, Sophia A. H. Heyde, Morten H. H. Nørholm

## Abstract

In *Escherichia coli*, one of the best understood microorganisms, much can still be learned about the basic interactions between transcription factors and promoters, particularly in the stationary phase. When a cAMP-deficient *cya* mutant is supplied with maltose as the main carbon source, mutations develop upstream from the two genes *malT* and *sdaC*. Here, we explore the regulation of the two promoters, using fluorescence-based genetic reporters in combination with both spontaneously evolved and systematically engineered *cis*-acting mutations. We show that in the *cya* mutant, regulation of *malT* and *sdaC* evolves toward cAMP-independence and increased expression in stationary phase. Furthermore, we show that the location of the cAMP receptor protein (Crp) binding site upstream of *malT* is important for alternative sigma factor usage. This provides new insights into the architecture of bacterial promoters and the global interplay between Crp and sigma factors in different growth phases.

## Introduction

The genetics of the model bacterium *Escherichia coli* is made up of hierarchical regulatory systems functioning on both local and global scales ^1,2^. On a local scale, transcription factors determine if conditions for transcriptional initiation are met, while global systems control entire genome subsets for example via sigma factors directing RNA polymerase (RNAP) to specific promoter motifs. Some transcription factors function on a global scale, such as the cyclic AMP receptor protein, Crp, which regulates the expression of hundreds of genes and acts like a nucleoid associated protein ^1,3,4^. In the absence of glucose, adenylate cyclase (Cya) produces cAMP, and Crp-cAMP then activates genetic programs that enable utilization of alternative carbon sources ^5^. Additionally, global regulation by Crp responds to deficiencies in nitrogen metabolism and synthesis of nucleic acids, making Crp a ubiquitous regulator for stress adaptation ^6,7^.

RpoS (σ^38^) is a main sigma factor in stationary phase, playing a role when cells have exhausted resources where growth stalls and bacterial ageing starts due to oxidative stress ^8–10^. Both Crp and RpoS are associated with starvation, but increased activity of Crp causes a decrease in intracellular RpoS, and correspondingly, strains with no Crp activity show increased resistance to oxidative stress due to increased levels of RpoS ^10,11^. Another sigma factor associated with stress is RpoH (σ^32^); classically termed a heat shock sigma factor, although knockouts are severely impaired already above room temperature ^12,13^ and a broader physiological role in stationary phase has been indicated ^14^. Several heat shock proteins are also important during starvation stress, and, thus, the heat shock implication and quasi-essentiality of RpoH may be due to its more general role in protein folding, for example in growth phase transitions where the cellular machinery goes through extensive reorganization ^14,15^.

Sigma factors of the RpoD family, which includes RpoS and RpoH, recognize specific promoter motifs termed the -35 and -10 boxes based on their positions in relation to the transcription start site (TSS, +1)^16^. In case of RpoD, the “house-keeping” sigma factor, the -35 box consensus sequence is 5’-T_35_T_34_G_33_A_32_C_31_A_30_-3’, and for the -10 box it is 5’-T_12_A_11_T_10_A_9_A_8_T_7_-3’. T_35_ and T_34_ are the most conserved, and A_11_ and T_7_ are the most important for open complex formation ^16–18^. Some promoters contain additional recognition sequences downstream (discriminator region, DR) and upstream (extended -10 boxes, ext-10) of the -10 box, depending on specific domains in the sigma factor. For example, the region 1.2 domain recognizes the DR ^19–21^.

The MalT transcriptional activator controls the synthesis of enzymes required for utilization of maltose and maltodextrins ^22^. Crp binds the P*malT* promoter at position -70.5 and is an example of a class I Crp promoter, where Crp binds upstream (centered at position -61.5, -71.5,-81.5 or -91.5) from the RNAP αCTD domain. This is opposed to class II Crp promoters, where Crp binding is centered at position -41.5 ^23–25^. The 10 bp-spacing regularity is due to the spatial alignment of Crp and the αCTD for every turn of the DNA helix ^23,26^.

In previous studies, using a cAMP-deficient *E. coli* strain, regions upstream from *malT* and *sdaC* were found to be mutational hotspots ^4,27^. Using fluorescence-based reporters of promoter activity, we here use these evolved *cis*-acting mutations as evolutionary clues to investigate the *malT* and *sdaC* promoter architectures in relation to different sigma factors and Crp and how gene expression adapts during long-term stress.

## Results

### Mutations develop upstream from *malT* and *sdaC* in starving bacterial colonies

Due to the absence of cAMP, the *cya* mutant of *Escherichia coli* K-12 MG1655 only forms small white colonies on maltose MacConkey agar, but upon prolonged incubation maltose fermenting (red) and non-fermenting (white) secondary colonies appear (Figure 1a) ^4,27^. Previously, in this evolutionary model system, mutations were identified upstream from *malT* ^7,27^ and *sdaC* ^4^: a P*malT\** mutation was isolated as a red secondary colony at 37°C, and two different P*sdaC*\* mutants were isolated as white secondary colonies both at 37°C and 44°C (Figure 1b and c). Looking at the DNA sequences, it appears that the three mutations result in putative -10 box sequences closer matching the RpoD consensus (Figure 1c). An identical P*malT\** mutation was isolated in another study and was shown to increase transcription of *malT* in the absence of Crp ^28^. This suggests that transcription initiation at these promoters is suboptimal under the conditions where the mutations developed, but does it reveal more about the specific promoter architectures?

**Figure 1.**
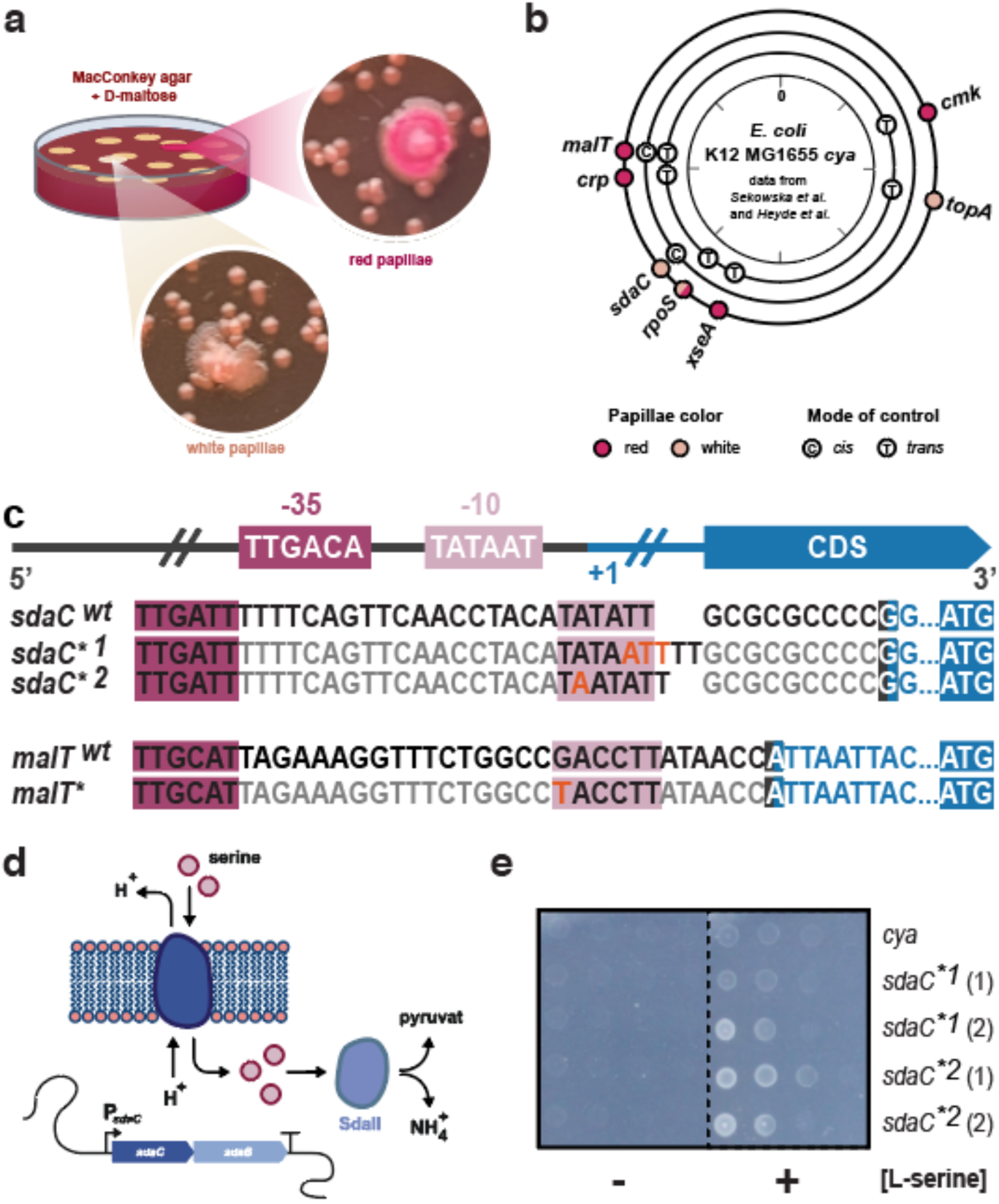
*malt* and *sdaC* mutations occurring in experimental evolution of *Escherichia coli*. a) Illustration of the experiment. *E. coli* K-12 MG1655 *cya* was plated on MacConkey maltose agar, small white colonies formed, and over time maltose fermenting (red) and non-fermenting (white) secondary colonies appeared. b) Mutational hotspots found previously ^4,7,27^ mapped by genomic locations, and marked by mutant phenotype and mutation type; red, fermenting; white, non-fermenting; (C), cis-acting mutation; (T), trans-acting mutation. c) Consensus sequence for RpoD, along with alignment of the *cis*-acting *malT* (SNP) and *sdaC* (insertions) promoter mutations (*) to their native (wt) sequences in *E. coli* K-12 MG1655. d) Illustration of the *sdaCB* operon and its function. L-serine import is mediated by the *sdaC*-encoded L-serine-H^+^-symporter, and degradation of L-serine to pyruvate and NH_4_^+^ is catalyzed by the L-serine deaminase SdaII. e) Serial dilutions of *E. coli* K-12 MG1655 *cya* with the native or mutant P*sdaC* promoters on M9 minimal media at 37°C in the presence or absence of L-serine as the sole carbon source. Two independent isolates of each mutation type were found and are marked (1) and (2).

### Mutations upstream from *sdaC* enable growth on L-serine

The *sdaC* promoter was previously shown to be regulated by Crp ^29^. The *sdaC* gene encodes a L- serine importer that enables the use of extracellular L-serine as an energy source (Figure 1d), and we speculated that the mutations in P*sdaC*\* could improve utilization of this amino acid. This was confirmed both at 37°C and 44°C when observing the growth of the P*sdaC*\* strains on minimal medium with L-serine as the sole carbon source (Figure 1e and Supplementary Figure 1).

### A fluorescent reporter predictably and sensitively reports on key regulatory features of P*malT*

In the *malT* promoter, only 3/6 and 2/6 nucleotides match the RpoD -35 box and -10 box consensus sequences, respectively (Figure 1c) ^28^. The A_11_ and T_7_ nucleotides important for open complex formation are conserved, and the P*malT*\* mutation adds the consensus T_12_, thereby potentially improving RpoD-mediated transcription, which could explain the selective advantage in a cAMP- deficient environment ^28,30^. On the other hand, RpoD should play only a minor role under the long-term starvation conditions where we isolated P*malT*\*, motivating further studies of the molecular mechanisms at play.

To further investigate the regulation of *malT*, we took advantage of a reporter plasmid, which harbors the superfolder *gfp* gene under control of P*malT* (Figure 2a) ^7^. The reporter is tightly repressed by glucose in *E. coli* K-12 MG1655 (Figure 2b), highly inducible and titratable by cAMP in a *cya* knockout strain (Figure 2c) ^7^, and sensitive to a cAMP-independent Crp* mutation ^7^ (Figure 2d). It therefore shows promise for providing insights into the dynamics of P*malT* regulation.

**Figure 2.**
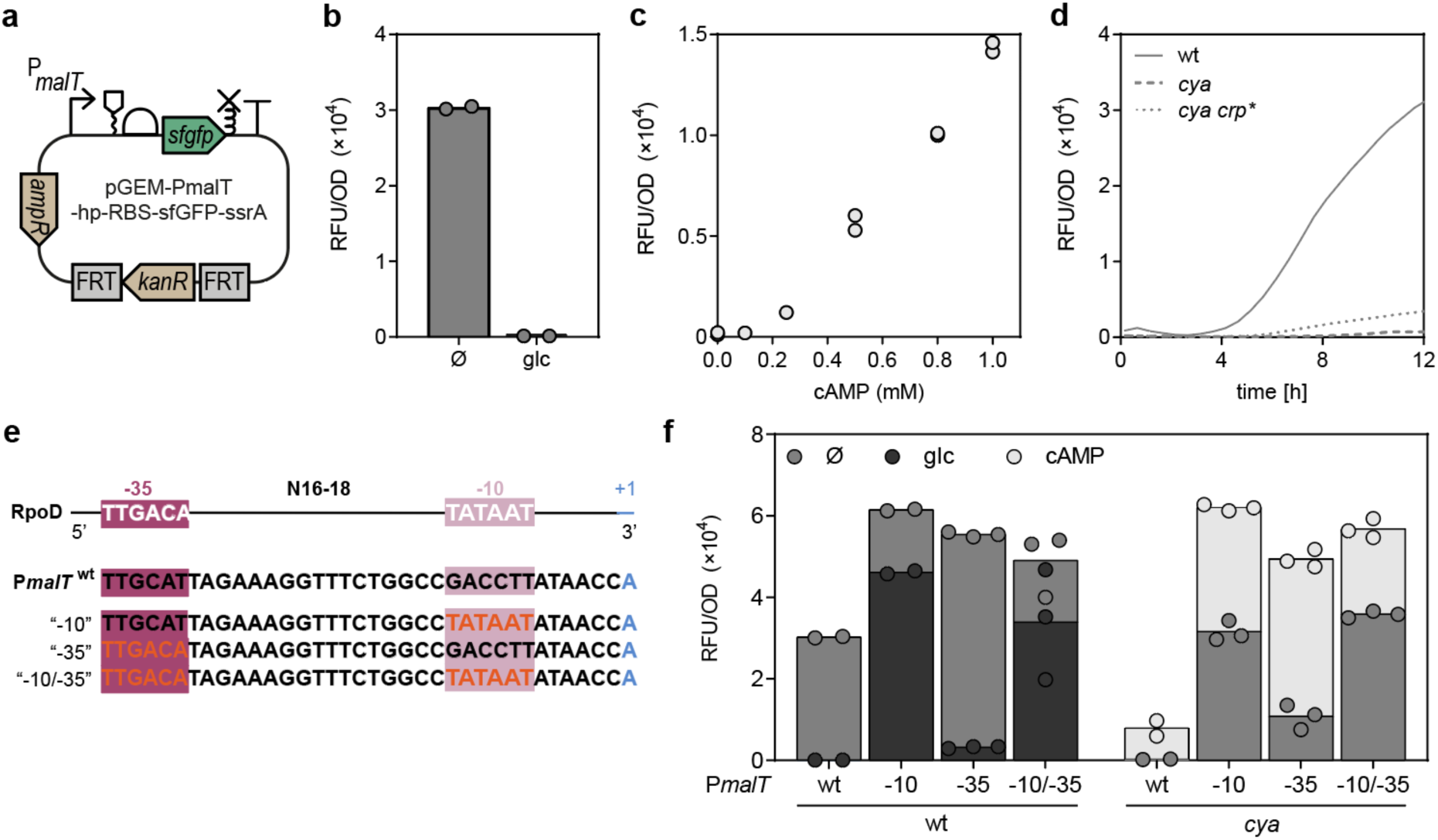
Characterization of the *malT* promoter and hybrid RpoD consensus mutants. a) Illustration of the reporter plasmid according to SBOL standards, elements being; a truncated *malT* promoter, without the Mlc site, an RNA-stability element, a ribosomal binding site, the coding sequence of superfolder GFP, an ssrA tag for protein degradation, and a terminator. The reporter also contains an ampicillin resistance marker for plasmid maintenance, and a kanamycin resistance marker with FRT sites for genomic integration. b) Fluorescence from the P*malT* reporter in *E. coli* K-12 MG1655 after 9 h of growth in the presence or absence (Ø) of glucose (glc, 1%). Data represents biological duplicates. c) Activity of the reporter in *E. coli* K-12 MG1655 *cya* after 9 h of growth with different levels of cAMP. Data represents biological duplicates. d) Fluorescence from the P*malT* reporter in *E. coli* K-12 MG1655, *cya*, and *cya crp\** through 12 h of growth. Data represents biological triplicates. e) Promoter layout of (top to bottom) RpoD consensus, the native P*malT* sequence from *E. coli* K-12 MG1655, and hybrid promoters mutants where the - 10 box, -35 box, or both are changed to the RpoD consensus (red font). f) Fluorescence from the native and consensus P*malT* reporter variants in *E. coli* K-12 MG1655 wt and *cya* after 12 h of growth, with no supplement (Ø, grey), 1% glucose (wt only, blue), or 0.5 mM cAMP (*cya* only, red). Data represents two or three biological replicates.

We first entertained a synthetic biology approach to better understand regulation of P*malT*. The poor similarity to the -10 box RpoD consensus sequence has been speculated to be the main cause of Crp-dependence ^30^, but the -35 box of P*malT* also differs from the RpoD consensus. This prompted us to systematically investigate changes in P*malT* towards the RpoD consensus. When converting either the -10 or -35 box to the exact RpoD consensus (Figure 2e), fluorescence from the P*malT* reporter increased as expected (Figure 2f). However, while the consensus -35 mutants retained almost full glucose repression and 4-fold cAMP induction, the -10 mutants only showed 20-30% glucose repression and 2-fold stimulation by cAMP (Figure 2f). This confirms that the low similarity of the -10 box to the RpoD consensus is the dominating cause of Crp-dependence_28_. Nevertheless, even for the exact RpoD consensus mutants (“-10/-35”) in the wild-type background, fluorescence was still stimulated by cAMP (+50%, Figure 2f).

### An alternative promoter may enable Crp-independent *malT* transcription in stationary phase

Looking at the *malT* promoter region, a close match for another -10 box (T_7_A_6_T_5_A_4_A_3_C_2_) is present five bp downstream from the previously annotated P*malT* -10 box (Figure 3a). We decided to investigate whether transcription from this alternative -10 box could be stimulated by re-aligning the Crp binding site position. To this end, we constructed mutants of the *malT* promoter region with the Crp binding site re-positioned from the natural -70.5 position to -75.5 (±1), -65.5 (±1), - 60.5, -55.5, and -50.5, and near class II positions 43.5 to -38.5 (Supplementary Figure 2). Fluorescence was monitored throughout growth, and is in Figure 3b plotted after 12 hours of growth, when the strain had entered stationary phase. However, the observed trends were consistent throughout all growth phases: In accordance with previous studies, local optima in promoter activity at sites corresponding to class I (-60.5) and class II (-42.5) Crp binding sites were observed (Figure 3b) ^23,24^. The lack of a local optimum at position -50.5 is expected due to the binding of the RNAP to the promoter at this position ^31^. Fluorescence as a result of a potential secondary transcript was significantly less intense than from the main transcript; we observed low signals from mutants -65.5 and -55.5, and thus, if a secondary promoter, as shown in Figure 3a, is indeed active *in vivo*, it is not stimulated by Crp binding at class I binding sites (Figure 3b).

**Figure 3.**
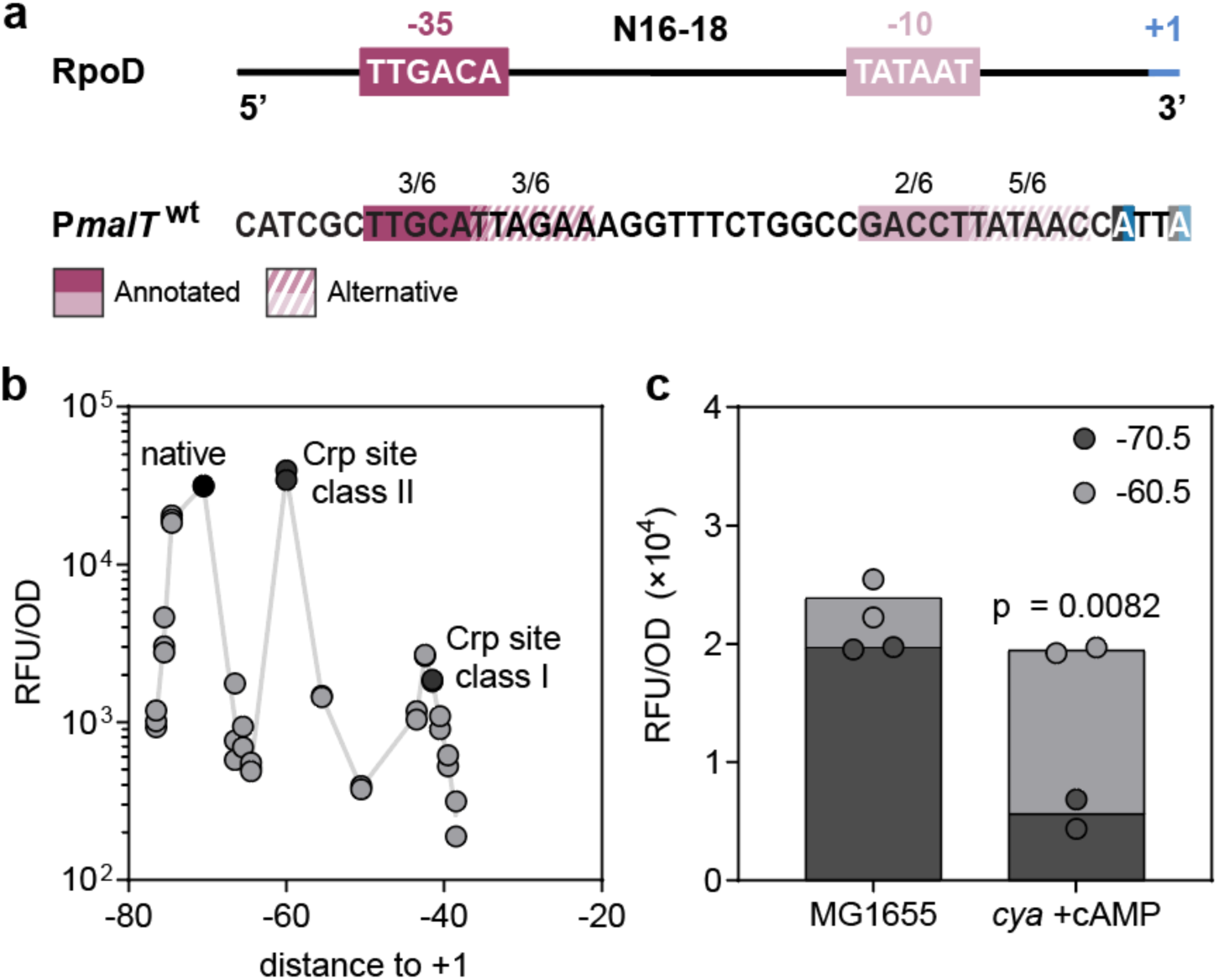
Effects of varying the Crp binding site location within the *malT* promoter. a) RpoD consensus sequences and the *malT* promoter with annotated and alternative RpoD recognition sequences and TSS - see main text for further description. The identity between the recognition sites and the consensus sequences are indicated with numbers above. b) Fluorescence from the P*malT* reporter variants after 12 h of growth. Indicated are the native distances between the Crp binding site and +1, and the distances classically associated with Crp promoter classes I and II. Data represents two or three biological replicates. c) Fluorescence from the native P*malT* reporter (-70.5) compared to the -60.5 variant in *E. coli* K-12 MG1655 WT or a *cya* strain supplemented with 0.5 mM cAMP after 12 h of growth. Significance was based on two-sided unpaired t-tests between the groups indicated and designated as significant if p < 0.05. Data represents biological duplicates.

### The dynamic range of expression is higher for a non-native P*malT* variant

An interesting observation, upon varying the distance between the annotated transcription start site and the Crp binding site, is that the fluorescence for P*malT* with Crp in position -60.5 is close to, or even higher, than that of the native promoter (Figure 3b), as previously observed ^23^. When looking at other RpoD promoters, activating Crp sites are more often annotated around position - 61.5 than at -71.5 (Supplementary Figure 3) ^32^; however, the evolutionary background for selection of one class I site position over others remains unelucidated. In the data shown in Figure 3c, the fluorescence for position -60.5 relative to position -70.5 was even higher when a Δ*cyaA* strain was supplemented with cAMP. Thus, in this assay, position -60.5 provides a larger dynamic range of expression than position -70.5. Nevertheless, the -70.5 position is preserved in K-12 MG1655. What might provide a selective advantage of binding Crp in this position? Do additional factors, besides those already known ^28,32–34^, play a role?

### Engineered RpoD consensus mutants show growth-phase-dependent effects

Given that different sigma factors dominate in different growth phases, we decided to look closer at the growth and time-resolved fluorescence of the RpoD consensus promoter mutants. As one would expect, all three P*malT* RpoD consensus mutants exhibited increased fluorescence compared to the wildtype P*malT* in the early stages of growth (Figure 4a, top). However, the two -10 box mutants exhibited an approximately two hours extended lag phase indicating early expression stress (Figure 4a). In the transition from exponential to stationary phase, none of the constructs exhibited an increase in fluorescence, showing either a temporary stall (-10 box mutants) or a local minimum. This “pause” before stationary phase could be due to changes in the transcriptional machinery involving sigma factors that take place around this point in growth ^35,36^. This, however, poses a new question: If RpoD is exchanged from the transcriptional machinery upon entry into stationary phase, exactly what sigma factor is responsible for the continued expression from P*malT* throughout stationary phase?

**Figure 4.**
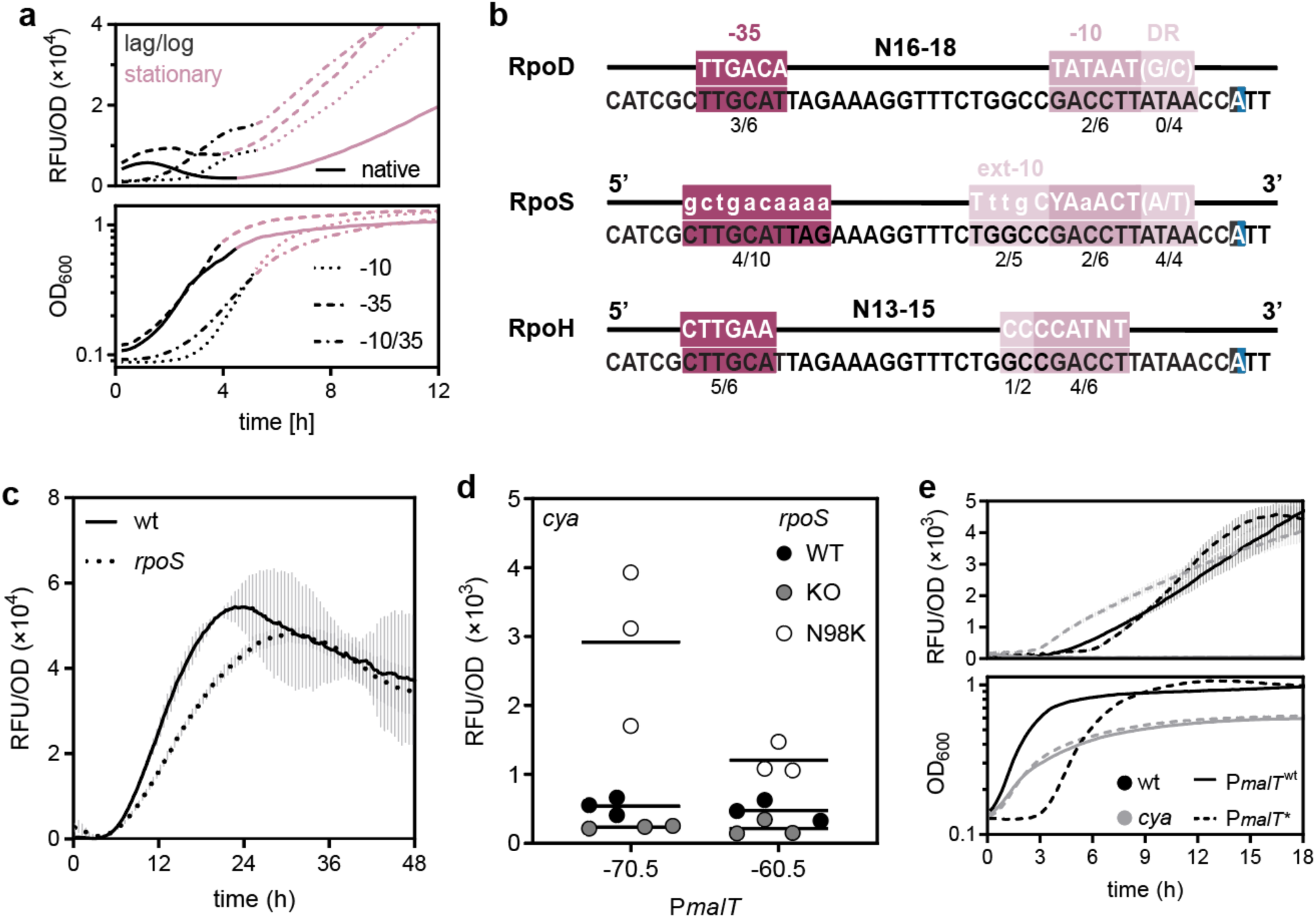
Expression from *malT* promoter reporters upon varying the central promoter motifs. a) Fluorescence (top) and growth (bottom) of *E. coli* K-12 MG1655 with the native or consensus P*malT* reporter variants during 12 h. Promoter variants: Solid line, native; dotted line, -10 consensus; stipled line, -35 consensus; dotted and stipled line, - 10 and 35 consensus. Black, lag and log phase; red, stationary phase. b) Alignment of P*malT* to the reported consensus sequences of RpoD, RpoS, and RpoH, with potential recognition sites. RpoS and RpoH both recognize extended -10 boxes, although it for RpoH often is designated part of the -10 box ^37^. c) Fluorescence with standard deviations from the P*malT* reporter in *E. coli* K-12 MG1655 wt and *rpoS* during 48 h of growth. Solid line, wt; stipled line, *rpoS*. d) Fluorescence from the native and -60.5 variant of P*malT* in *E. coli* K-12 MG1655 *cya crp* pSEVA-*crp* with various *rpoS* genotypes after 12 h of growth. Grey, *rpoS^+^*; red, *rpoS^-^*; blue, *rpoS* N98K. e) Fluorescence with standard deviations (top) and growth (bottom) of *E. coli* K-12 MG1655 with the P*malT*\* reporter during 18 h. Solid line, native promoter; stipled line, evolved promoter; black lines, wild-type strain; grey lines, *cya*. Data represents biological triplicates.

### The *malT* promoter shows similarities to alternative sigma factor recognition sites

Both RpoS and RpoH are associated with stress responses and stationary phase ^14,36,38^ and could be involved in *malT* expression. The DNA sequence of P*malT* is a potential good match for RpoS, when taking into account the nucleotides in the RpoS ext-10 box (T_17_ and C_13_), and the discriminator region, which is A-T rich, as reported optimal for RpoS (Figure 4b) ^39^. However, motifs can be found that match the RpoH consensus as well (Figure 4b) ^7,37,40^. It is also worth noting that the P*malT* mutants with an RpoD consensus -10 box retain similarity to the RpoS but not the RpoH consensus sequence. Thus, one hypothesis for the observed Crp-independence of these mutants could be that an RpoD-consensus -10 box causes a shift away from RpoH-driven transcription towards RpoS-driven transcription.

### The *malT* promoter is affected by different *rpoS* mutants

Potential effects of RpoS on P*malT* were explored by monitoring the activity of the native *malT* promoter in mutant *rpoS* strains. Compared to the *rpoS+* control, an *rpoS* knockout strain shows a significant decrease in fluorescence in early stationary phase (Figure 4c). This indicates involvement of RpoS in expression of *malT*.

Crp-independent effects were assayed in the *cya* mutant background where low levels of fluorescence were observed after 12 h, both in the presence and absence of *rpoS*. An RpoS_N98K mutant evolved on several independent occasions during long-term starvation in a previous study _27_ and this mutant showed increased fluorescent levels from the *malT* promoter (Figure 4d), but not the synthetic -60.5 P*malT* variant. This provides independent evidence for RpoS-driven transcription and suggests that it is dependent on upstream elements that are compromised in the - 60.5 mutant (Figure 4d). Unfortunately, experiments to test potential *in vivo* direct interaction between RpoH and P*malT* were inconclusive due to the severe physiological changes caused by *rpoH* deletion.

### The P*malT\** mutation enables *malT* expression in the absence of cAMP

The potential interplay between a stationary phase sigma factor and P*malT* is not in conflict with the naturally evolved *cis*-acting mutation in P*malT* (Figure 1c), as the similarity to the consensus sequences are preserved for RpoH and improved for RpoS. To further explore the consequence of the P*malT*\* mutation, we introduced the point mutation into the *gfp* reporter construct. Similar to the -10 box mutants, when expressing *gfp* from P*malT*\* in a wild-type strain, growth exhibits a 3 hour extended lag phase, indicating expression stress (Figure 4e). However, the clearest effect was seen in the difference in fluorescence between P*malT* and P*malT*\* in a *cya* strain, where fluorescence was completely absent from P*malT* but high with P*malT*\* (Figure 4e). This clearly explains the selective advantage of the P*malT*\* mutation for growth on maltose in a *cya* strain.

### *sdaC* is mainly expressed in stationary-phase and is increased in *sdaC\** mutants

Returning to *sdaC,* as interactions of alternative sigma factors with P*malT* are indicated in the data presented here, it is not unreasonable to suspect that the *cis*-acting mutations that evolved in P*sdaC* under similar conditions similarly relate to sigma factor usage. Except for binding sites for Crp and Lrp ^29,41^, only few *sdaC* promoter features have been described. A TSS for an RpoD promoter has been annotated, and an RpoH-driven promoter has previously been implicated upstream from *sdaC*, but no promoter has been experimentally verified ^42,43^. Similar to *malT*, we monitored the growth and time-resolved fluorescence from P*sdaC*-GFP reporters with or without the P*sdaC*\* mutations (Figure 1c). The mutant promoters showed increased fluorescence from early stationary phase, with an additional increase occurring after additional ∼3-4 hours in stationary phase (Figure 5a). At 44°C, the temperature where some of the P*sdaC*\* mutants were isolated, the P*sdaC*\* mutants show consistently higher levels of fluorescence compared to the native promoter in both exponential and stationary phase (Supplementary Figure 1). To test the involvement of RpoS in regulation of P*sdaC*, we assayed the wild-type and mutant promoters in an *rpoS* background. The two mutant promoters, especially P*sdaC*\**2*, showed decreased fluorescence in the *rpoS* knockout background, while the wild-type promoter was unaffected (Figure 5b). This implicates involvement of RpoS in regulation of the mutant *sdaC* promoters.

**Figure 5.**
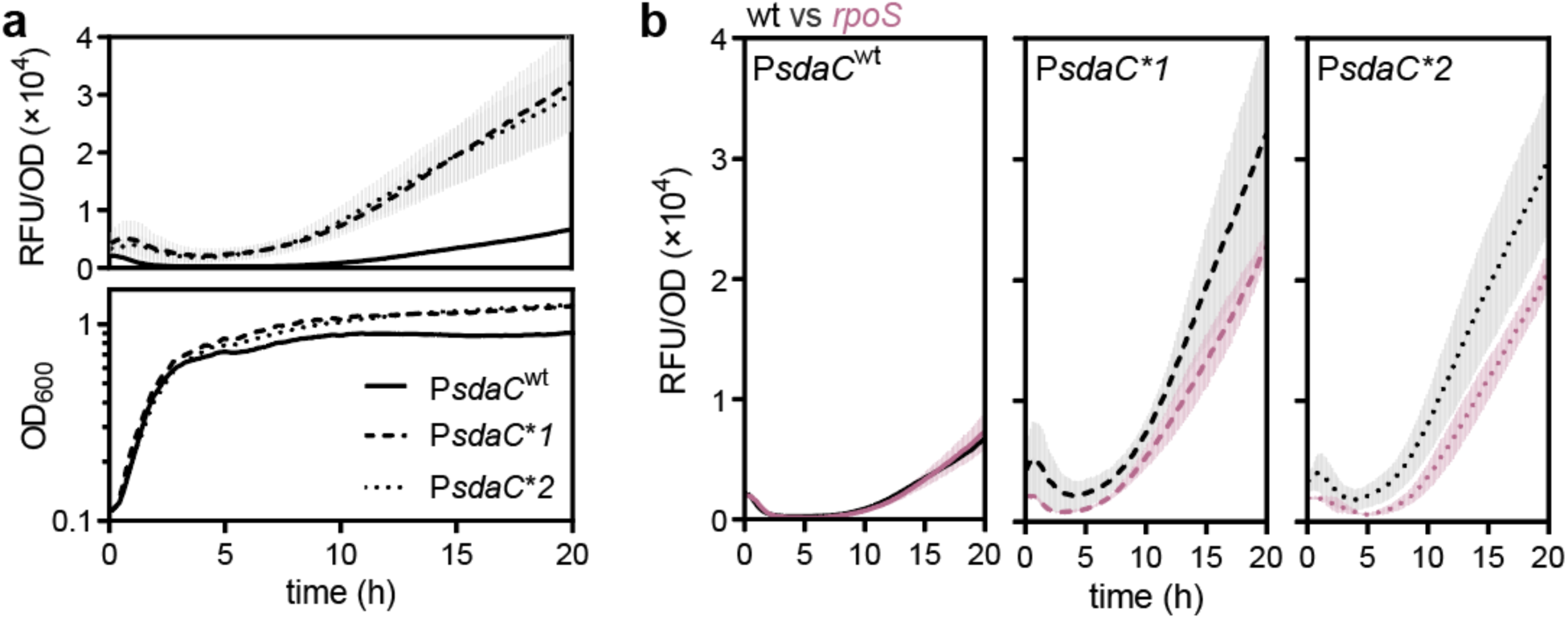
Growth-phase dependent regulation of P*sdaC*. a) Fluorescence with standard deviation (top) during 20 hours of growth (bottom) of *E. coli* K-12 MG1655 with the P*sdaC* reporter mutants. Promoter variants: Solid line, native (wt); stipled line, mutant 1; dotted line, mutant 2. b) Fluorescence with standard deviations from the native P*sdaC* (left), P*sdaC*\**1* (middle), and P*sdaC*\**2* (right) in *E. coli* K-12 MG1655 wt (grey) and *rpoS* (red) through 20 h of growth. Solid line, native (wt); stipled line, mutant 1; dotted line, mutant 2.

## Discussion

Promoter-activating mutations are frequent outcomes of evolution ^44^, and have led to important insights into the mechanisms of regulation ^45^. The *cis-*acting mutations characterized here explain well the growth benefits gained under the specific selective conditions: As P*malT*\* was isolated at 37°C on maltose, the main selection regime was carbon starvation, which would be abolished, should the cell gain the ability to utilize maltose; a likely outcome of increased *malT* expression. Similarly, P*sdaC* controls the *sdaCB* operon, and overexpression may result in an increased rate of L-serine import into the cell and the use of L-serine in an energy-providing role ^46^. Selection of P*sdaC*\* at 37°C and 44°C thus implies that L-serine can be used as a carbon source from MacConkey agar during both starvation and heat stress. Alternatively, there is strong selection pressure to express *sdaC* to prevent cell lysis during starvation ^47^.

In both cases mentioned above, it can be reasoned that the relieve of stress is achieved via adapting the promoter in question to the transcriptional machinery active at the time of the mutational event. The mutations were isolated following days or weeks of starvation stress (and for some P*sdaC* mutations, additional heat stress), causing growth to stall. Since RpoD is mostly unavailable under these conditions, and as RpoS is stabilized during conditions such as long-term starvation and heat stress, it seems unlikely that the mutants were rescued from stress based on *cis*- acting mutations improving RpoD interactions ^10,15,35^.

The *sdaC* promoter, as annotated in this study, shows a great match to the RpoD consensus, especially when considering the high GC content in the discriminator region and the lack of an extended -10 sequence. RpoD and RpoS consensus sequences show significant overlap, and the difference in transcription of some RpoD/RpoS promoters may simply be growth phase dependent, as RpoS shows a higher affinity for relaxed DNA in stationary phase ^48^. Insertions of +ATT (*\*1*) or +A(*\*2*) in the -10 box of P*sdaC* elongates the AT-rich sequence of this promoter, resulting in a discriminator region with diminished GC content, which may improve RpoS recognition of the promoter. Indeed, P*sdaC*\**1/2* show increased promoter activity through both exponential and stationary phase, and a knock-out of *rpoS* diminishes this effect somewhat, indicating dual transcription by RpoD and the stationary phase sigma factor. Similarly, the mutation in P*malT*\*, while changing the consensus of the -10 box towards the RpoD consensus, simultaneously improves the similarity to the RpoS consensus. When considering the already AT-rich discriminator region, and a match for a potential extended -10 region, the native *malT* promoter, while annotated as RpoD-transcribed, may very well be transcribed by RpoS as well; a hypothesis supported by the *in vivo* data presented here. The native *malT* promoter also contains motifs that may indicate RpoH recognition, and the P*malT*\* mutations were previously isolated during long-term starvation ^28,30^. Thus, it seems that there is a connection between P*malT*\* and the expression context that is long-term stationary phase, suggesting that P*malT*\* gains the ability to be transcribed by RpoS or RpoH, while native RpoD-dependent and possibly RpoH-dependent transcriptional programs are preserved.

As RpoS partially controls transcription of *rpoH* in stationary phase, the RpoS-dependent increased transcription from P*malT*\* and P*sdaC*\**1/2* could implicate either sigma factor. The high frequency of *rpoS* knockout mutations isolated during long-term starvation at both 37°C and 44°C _4,27_, and a mutation found in *ftsH* ^4^, a protease that degrades RpoH, could suggest that although RpoS may recognize the promoters, expression using RpoH is more important for long-term survival. The interplay between RpoS and RpoH should also be considered in the context of the frequently observed *rpoS* knockout mutations that are associated with a growth advantage in stationary phase ^49,50^.

Besides its status as heat shock sigma factor, RpoH is implicated in general stress responses as a key regulator of chaperones and proteases, ensuring homeostasis of protein function in *E. coli*. RpoH is also, directly or indirectly, implicated in starvation responses ^14,51^, DNA relaxation (independently from heat-shock induced DNA relaxation) ^52^, and oxidative stress resistance in exponential phase ^53^. RpoH has previously been suspected to be involved in regulation of the *malT* promoter; in previous work, the supplementation of cytidine to a strain carrying the P*malT* reporter caused a reduction in fluorescence output, while P*malT*\* abolished this effect ^7^. Cytidine binds to CytR, which represses *rpoH* transcription, releasing repression and potentially resulting in increased RpoH levels. P*malT* activity could thus be diminished in the presence of cytidine due to decreased availability of RNAP-RpoD relative to RNAP-RpoH; however, the lack of a cytidine response for P*malT*\* rather suggests that the repressive effect of cytidine on P*malT* is mediated by the sigma factor recognition of P*malT*, most likely via RpoH. Due to the difficulties in assaying RpoH effects on promoter activity without perturbing the entire genetic system by low growth temperatures, RpoH interactions with the *malT* promoter need further investigation.

Considering how the consensus sequences for RpoD and RpoH partially overlap and can coexist in one promoter, it is possible that during certain environmental conditions or metabolic states, RpoH can transcribe promoters otherwise designated RpoD, e.g., a hypothetical scenario during carbon starvation when RpoD is sequestered and RpoS levels are decreased due to Crp activity. RpoH is known to undergo structural changes upon induction of the heat shock response, so promoter recognition may even change depending on the temperature. One can imagine how functional overlap of the transcription machinery may be beneficial for the fitness and adaptability of evolved microbes; it allows the cell to rapidly respond to changes in environmental conditions and survive until the transcription machinery is exchanged in favor of long-term stress responses. The observations reported here indicate that P*malT* and P*sdaC*\* are recognizable by RpoS, and potentially RpoH, although RpoD-dependent transcription may be favored in the presence of Crp-cAMP, and hints that this interplay is affected by the location of the Crp-binding site in a class I Crp promoter. Thus, the regulation of the *malT* operon, and other metabolic genes, is based on an intricate interplay between Crp and sigma factors depending on the metabolic state.

## Acknowledgements

This work was supported by a grant from the Novo Nordisk Foundation NNF17OC0027752 to MHHN.

## Supplementary materials for

## Methods

### Bacterial strains

All strains applied were *E. coli* K-12 MG1655. The *cya* deletion (*cya::cat*) was introduced into a Δ*fnr* background by λRed recombineering in previous work ^1^. Experimental evolution of *cis*-acting mutations, the *crp*\* (*crp*A144T), as well as *rpoS*N98K, in the *cya* background were described previously ^1,2^. The *rpoS* gene was deleted in wild-type and *cya* backgrounds using *λ*Red recombineering with pSIM19 as previously described ^3,4^ and integration of a *tetA* PCR product with homology ends to the *rpoS* locus as previously described ^5^. In order to remove and avoid secondary mutations in *crp*, the *crp* gene was further deleted in *cya rpoS*N98K, *cya rpoS*, and *cya*, and supplied on a low-copy plasmid described previously ^6^.

### Growth conditions

Strains were grown in LB broth or on LB agar at 37°C unless otherwise mentioned and supplemented with ampicillin (100 μg/ml) or kanamycin (50 μg/ml) for plasmid maintenance when necessary. For characterization of serine utilization in *sdaC*\* mutants, strains were drop-tested on M9 minimal medium supplemented with 2% L-serine. Supplements used for growth experiments were additionally cAMP (0.5 mM) or glucose (1%) as denoted.

### Plasmid construction

Plasmids were constructed by USER cloning of relevant fragments ^7^ and sequenced before application. All plasmids are shown in Supplementary Table 1 and oligonucleotides used for plasmid constructions as listed below are shown in Supplementary Table 2. The P*malT* reporter variants were constructed as one-fragment clonings based on the native P*malT* reporter as follows: Variants -76.5 to -64.5 were constructed using oligo #5733 in combination with oligos #5727- #5732, variants -60.5 to -38.5 were constructed using oligo #5590 in combination with oligos #5591-#5600, and variants -10, -35, and -10/-35 were constructed using oligo #5726 in combination with oligos #5734-#5736. The P*sdaC* reporters were constructed based on the native P*malT* reporter as a vector backbone, using oligos #4256 and #4258, and the genomic *sdaC* promoter amplified from the native or mutant strains using oligos #4577 and #4578.

### Growth experiments

Fresh colonies from cryostock were inoculated into a 5 mL preculture and grown over day (7 hours). From the preculture, 1 µL was used for inoculation into 150 µL medium in a 96-well microtiter plate (Costar 96 flat bottom), which was sealed using a breath-seal film and incubated in a Synergy H1 microplate reader (BioTek) (20-48 h, 37*°*C or 44*°*C, orbital shaking continuously at 425 cpm, reading depth of 3 mm). Optical density at 600 nm and fluorescence (excitation 485 nm, emission 528 nm, gain 50–100) was measured every 10-15 minutes. Data output in Microsoft Excel was subjected to preprocessing and visualized and analyzed using GraphPad Prism 9.1.0.

### Prediction of Crp binding sites

The PredCRP program was downloaded from github and executed in the Unix terminal using Python. The intergenic region between *ppnN* and *sdaC* was used as input (reference sequence NC_000913.3). The sequence was evaluated using a 42-bp moving window, and plotted using the position of the base in the center right of the moving window. The output of PredCRP was exported as .csv and is included in Microsoft Excel format as Supplementary Data 2.

## Data availability

Source data for this paper are available from the authors.

---

**Supplementary Figure 1.**
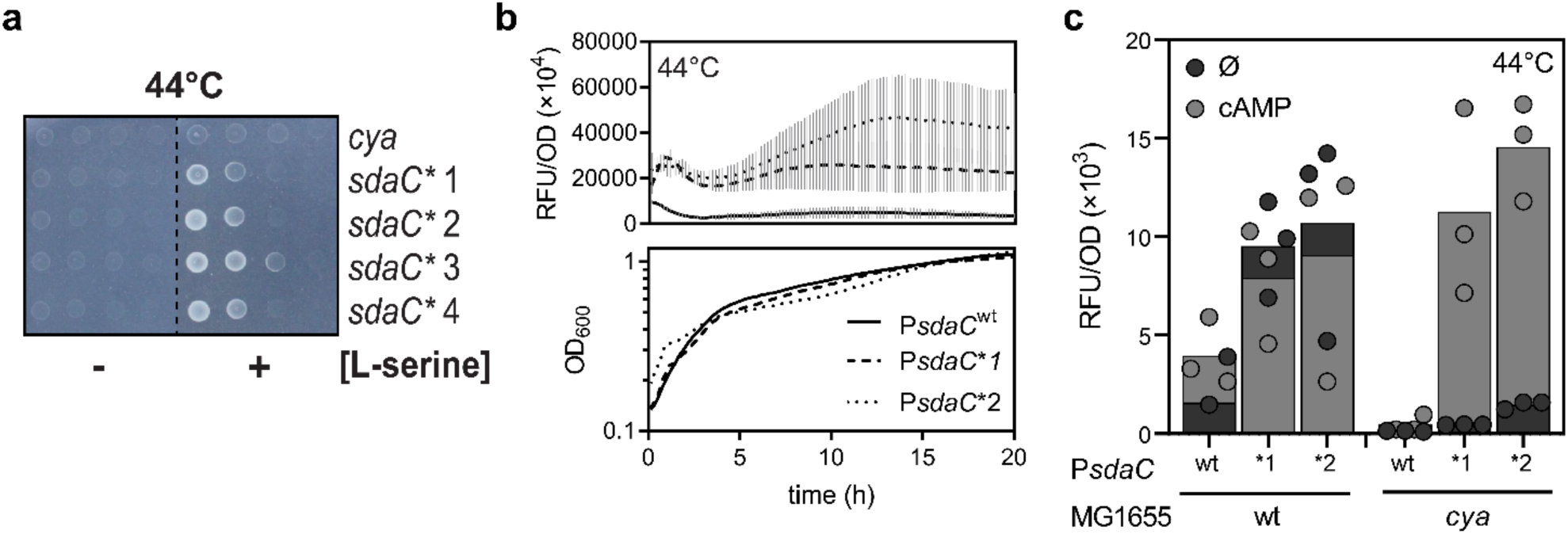
Effects of P*sdaC*\* at 44*°*C. a) Serial dilutions of *E. coli* K-12 MG1655 *cya* with the native or one of two promoter mutations in P*sdaC*, on M9 minimal media at 37*°*C in the presence or absence of serine as a carbon source. Two mutants of each mutation type were found and are marked (1) and (2). b) Promoter activity for the native (wt) and mutant (*) *sdaC* promoters in *E. coli* K-12 wild-type and *cya* after 20 h of growth at 44*°*C. For each strain-reporter combination, two growth conditions are superimposed; growth with no supplement (Ø, dark grey) or in the presence of 0.5 mM cAMP (grey). c) Fluorescence from the native (wt) and mutant (*) *sdaC* promoters in *E. coli* K-12 wild-type and *cya* after 12 h of growth at 44*°*C. For each strain-reporter combination, two growth conditions are superimposed (not staggered); growth with no supplement (Ø, dark grey) or in the presence of 0.5 mM cAMP (grey). The tallest bar is shown in the back, and the shortest in front.

**Supplementary Figure 2.**
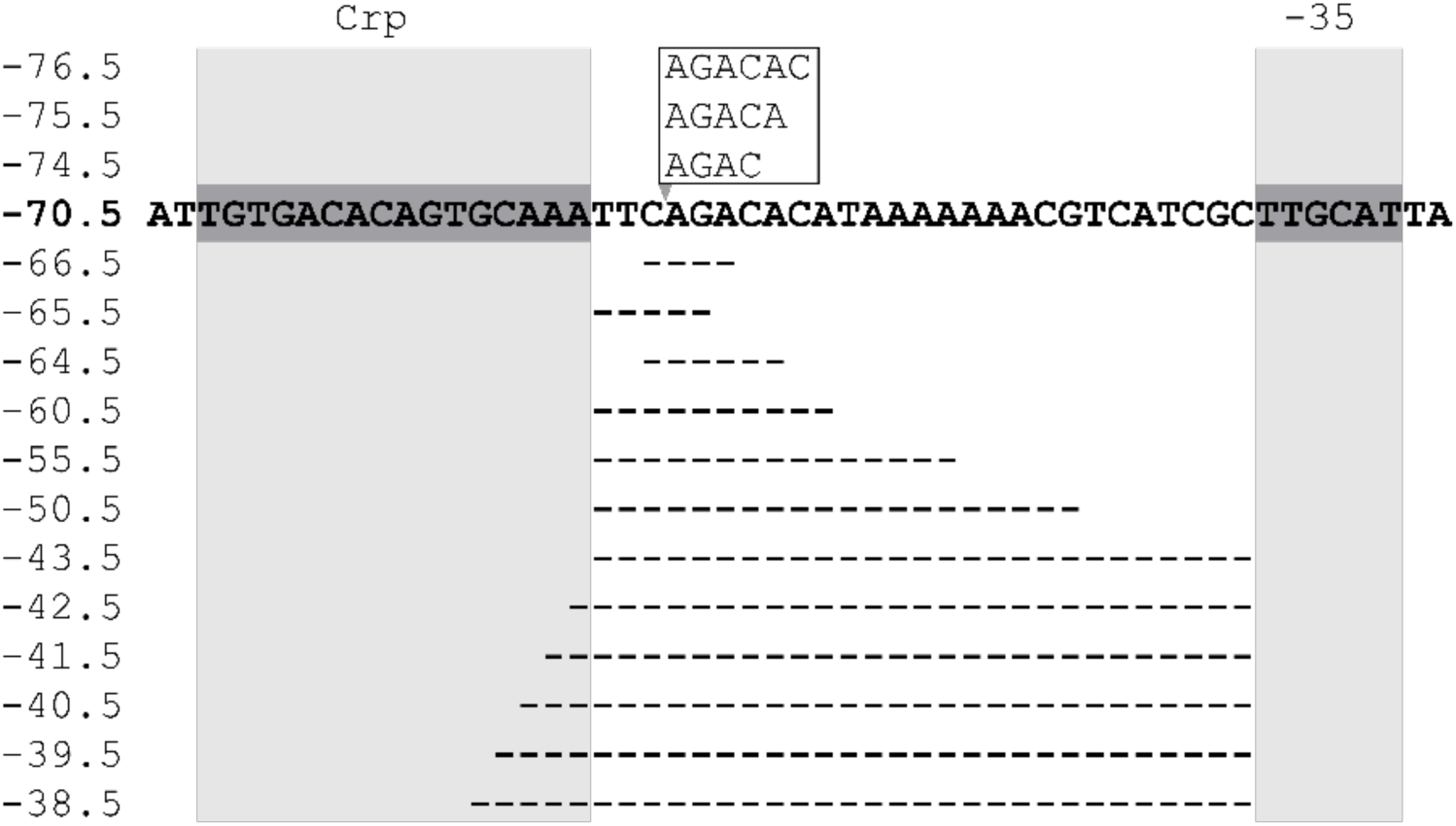
The native *malT* promoter and the Crp distance mutants investigated in this study.

**Supplementary Figure 3.**
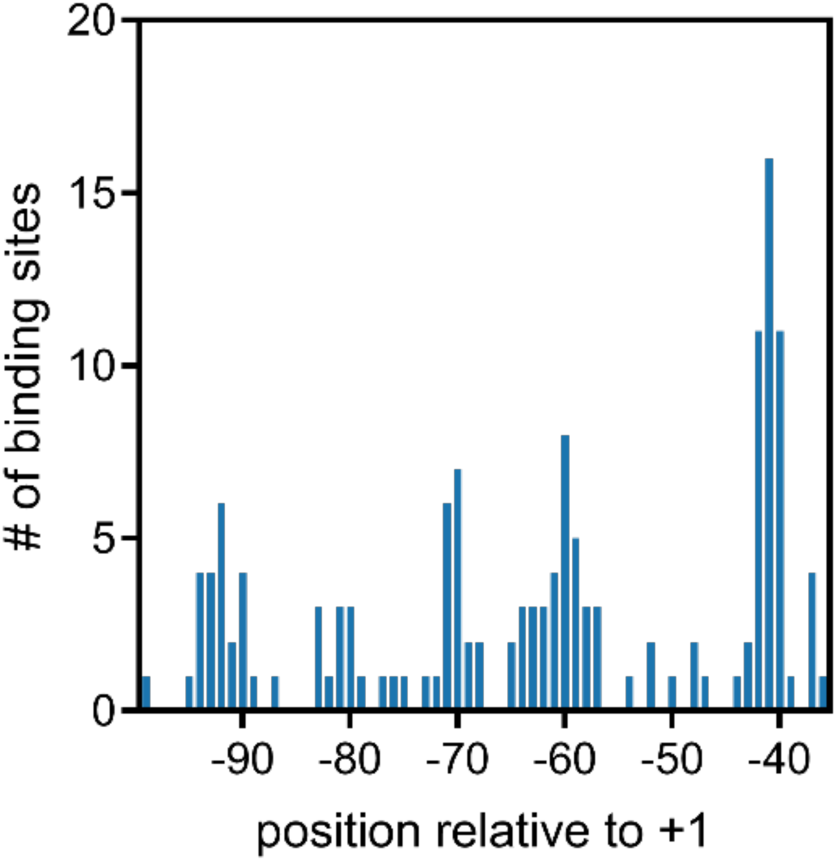
Distribution of activating Crp binding sites on RpoD promoters. Data from RegulonDB, 26-11-2021 (Santos-Zavaleta 2019).

**Supplementary Table 1.**
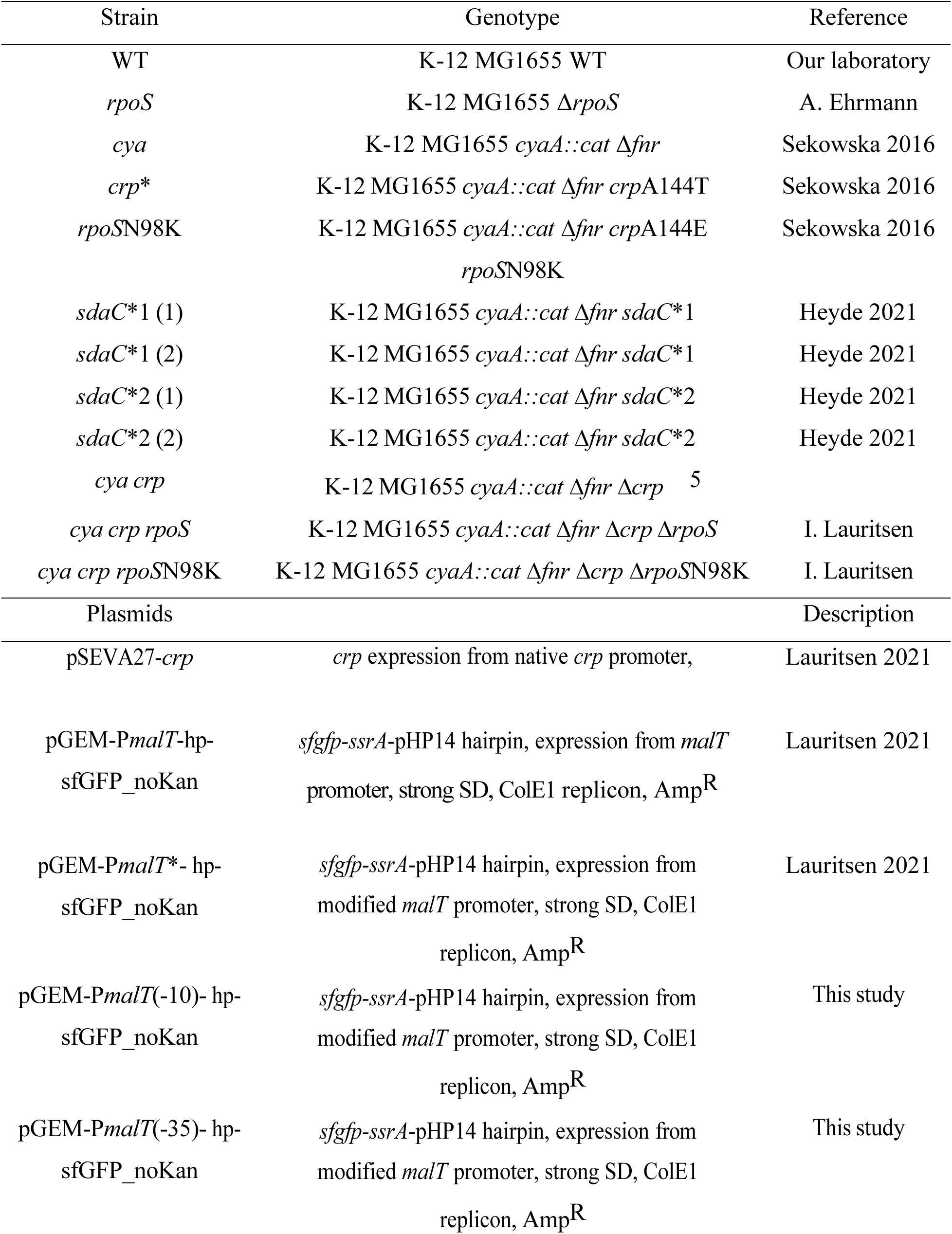
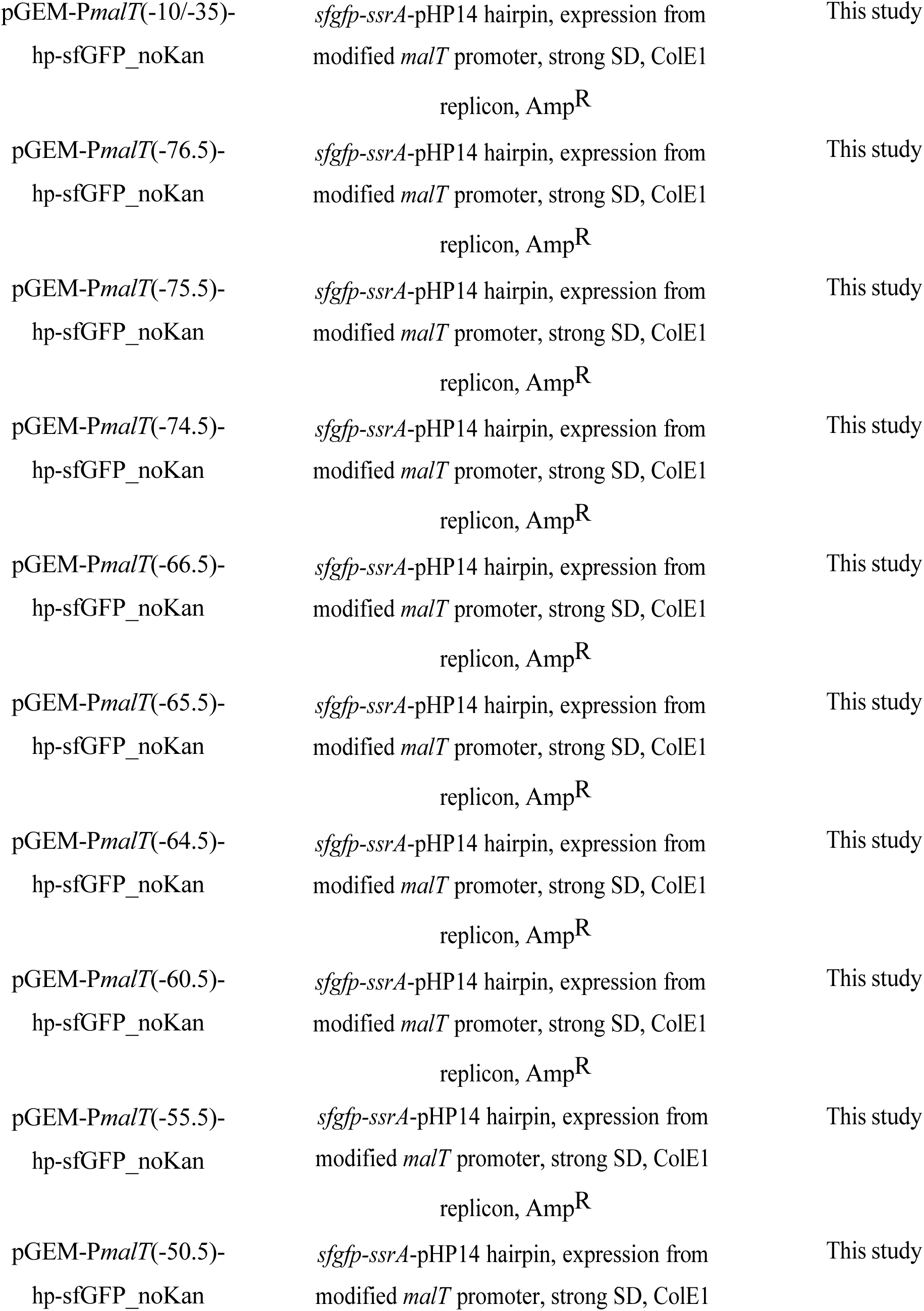
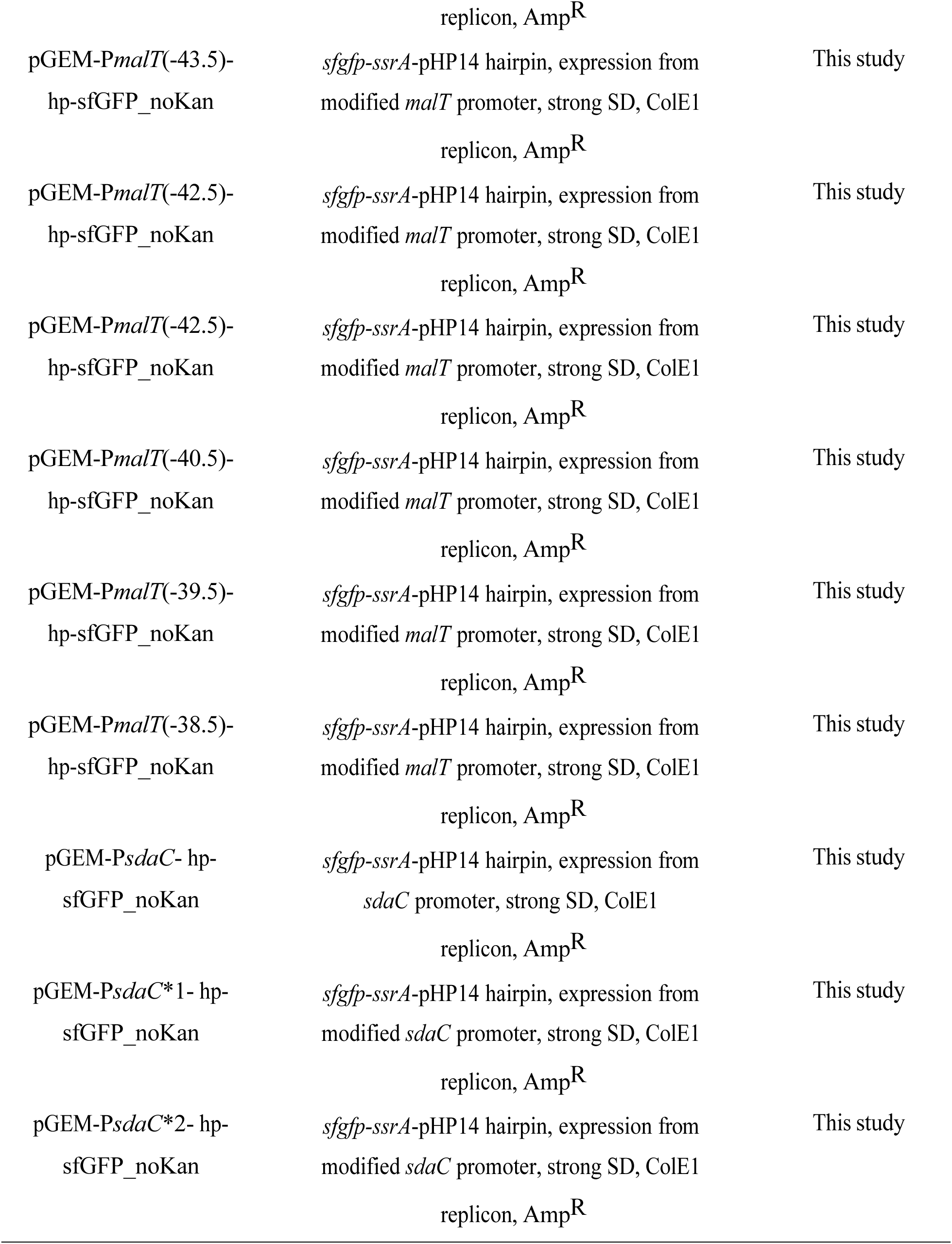
Strains and plasmids used in this study.

**Supplementary Table 2.**
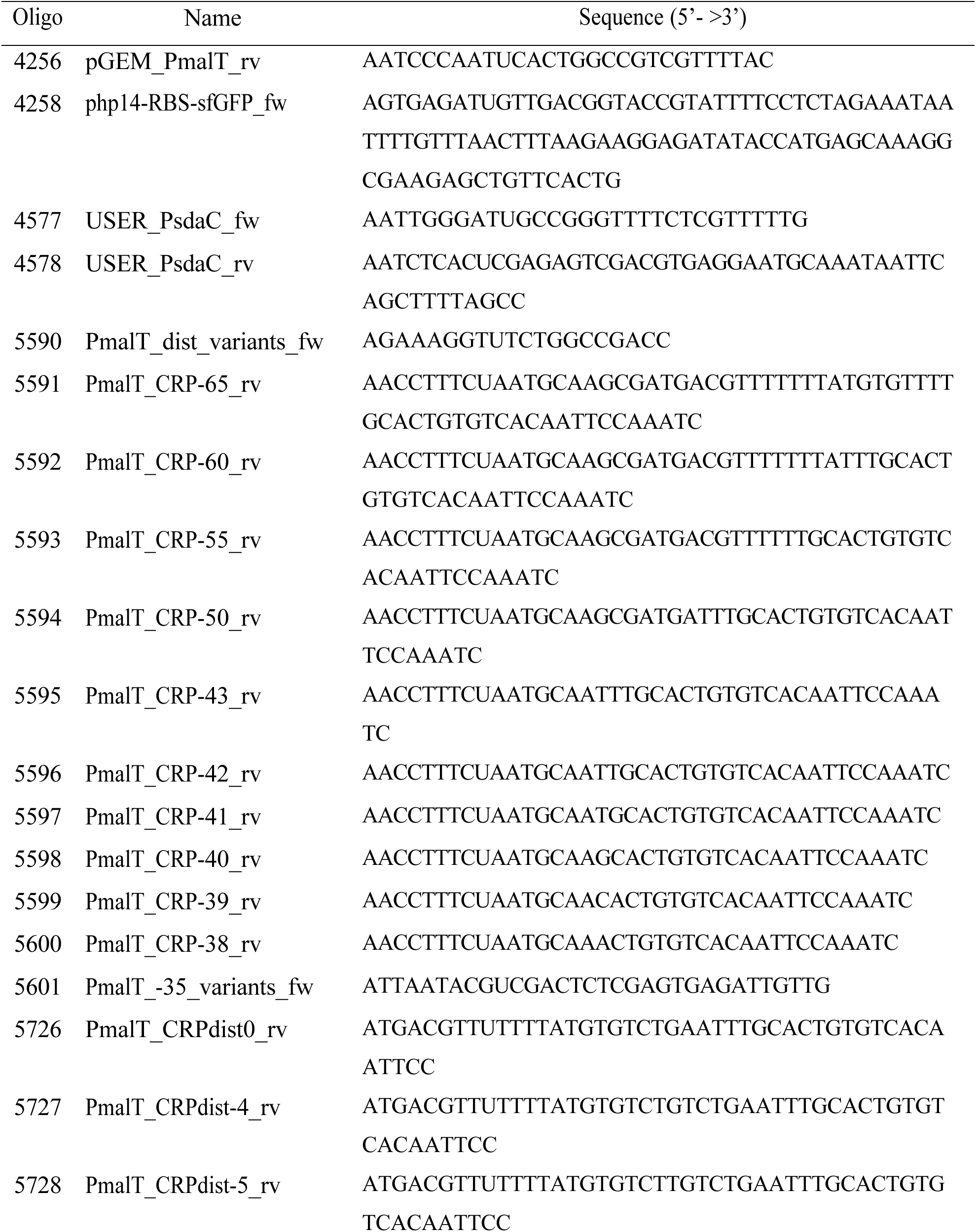
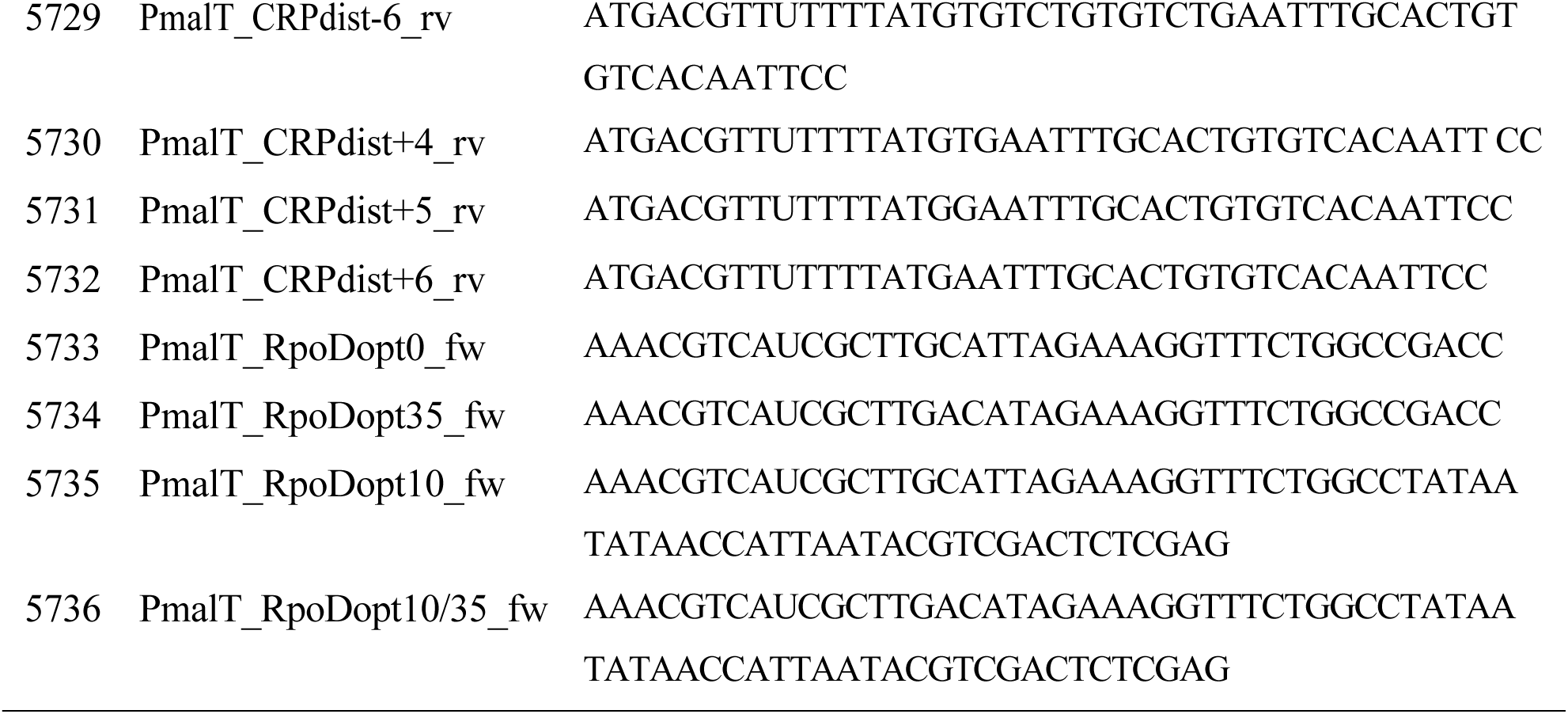
Oligonucleotides used in this study.

